# Does rhythmic priming improve grammatical processing in Hungarian-speaking children with and without Developmental Language Disorder?

**DOI:** 10.1101/2020.06.19.162347

**Authors:** Enikő Ladányi, Ágnes Lukács, Judit Gervain

**Author notes:** Corresponding author: Enikő Ladányi, 1215 21st Avenue South, 10420B Medical Center East, South Tower, Nashville, TN 37232.

## Abstract

Research has described several features shared between musical rhythm and speech or language, and experimental studies consistently show associations between performance on tasks in the two domains as well as impaired rhythm processing in children with language disorders. Motivated by these results, in the current study our first aim was to explore whether the activation of the shared system underlying rhythm and language processing with a regular musical rhythm can improve subsequent grammatical processing in preschool-aged Hungarian-speaking children with and without Developmental Language Disorder (DLD). Second, we investigated whether rhythmic priming is specific to grammar processing by assessing priming in two additional domains: a linguistic but non-grammatical task (picture naming) and a non-linguistic task (nonverbal Stroop task). Third, to confirm that the rhythmic priming effect originates from the facilitating effect of the regular rhythm and not the negative effect of the control condition, we added a third condition, silence, for all the three tasks. Both groups of children showed better performance on the grammaticality judgment task in the regular compared to both the irregular and the silent conditions but no such effect appeared in the non-grammatical and non-linguistic tasks. These results suggest that 1) rhythmic priming can improve grammatical processing in Hungarian, a language with complex morphosyntax, both in children with and without DLD, 2) the effect is specific to grammar and 3) is a result of the facilitating effect of the regular rhythm.

**Research Highlights:**

- 6-year-old Hungarian-speaking children with and without Developmental Language Disorder perform better on a grammatical task subsequent to exposure to a regular rhythm vs. an irregular rhythm/silence
- The effect of regular rhythm is specific: it improves performance on a grammatical task but not on a word retrieval or a non-linguistic task
- Difference between performance following regular vs. irregular rhythm originates from the facilitating effect of the regular rhythm (not the negative effect of the irregular rhythm)
- The results highlight the importance of rhythm in speech processing, and point towards a possible intervention tool in language disorders

## 1. Introduction

### 1.1 Rhythm in language

In our everyday life, we continuously produce and process different rhythmic stimuli. In some cases, the role of rhythm is more evident, such as when we listen to or play music, while in other cases, it is less outstanding such as when we chew. Rhythm seems to play a crucial role in speech production and processing as well. Indeed, languages have their characteristic rhythms. Linguists have long established (Abercrombie, 1967; James 1940; Ladefoged, 1993; Pike, 1945) that the languages of the world may be grouped into three categories on the basis of their rhythmicity, traditionally categorized as syllable-timed languages, such as Spanish or Italian, stress-timed languages, like English, Dutch or Arabic, and mora-timed languages, like Japanese and Tamil (Ramus, Nespor & Mehler, 1999).

These differences are easy to perceive. English and Dutch sound like Morse code, with consonants tending to cluster together (e.g., English *springs*), while Spanish or French have what is sometimes referred to as a machine-gun rhythm, with a more equally paced alternation of consonants and vowels (e.g., Spanish *bufanda* ‘scarf’). We rapidly realize if two rhythmically different languages are spoken in our environment, even if we never heard either of the languages before. Rhythmic differences across languages are actually so salient that even newborn infants and monkeys are also able to perceive them (Mehler, Jusczyk, Lambertz, Halsted, Bertoncini, & Amiel-Tison, 1988; Ramus, Hauser, Miller, Morris, & Mehler, 2000). Linguistically, rhythmic differences between languages have been linked to the relative proportions of vowels and consonants and their distribution in speech (Dellwo, 2006; Grabe & Low, 2002; Loukina, Kochanski, Rosner, Keane, & Shih, 2011; Ramus et al., 1999; Wiget, White, Schuppler, Grenon, Rauch, & Mattys, 2010). Acoustically, they are captured by the amplitude modulation rate of the auditory speech signal in different languages (Varnet, Ortiz-Barajas, Guevara Erra, Gervain, & Lorenzi, 2017).

Underlying the strong rhythmicity of the speech signal, linguistic constituents making up speech follow each other at a regular pace at multiple levels (Langus, Mehler, & Nespor, 2017). Phonemic changes such as formant transitions (differentiating /ba/ and /da/) or the coding of voicing (differentiating /ba/ and /pa/) occur at a high modulation frequency (~30-50 Hz). Syllables follow each other at a 4-7 Hz rate, while lexical and phrasal units occur at a 1-2 Hz modulation rate. Although the intervals between these units are not completely isochronous, they are rhythmic enough to elicit robust regularities in the time domain of speech (Giraud & Poeppel, 2012). Several studies indicate the importance of the regular timing of stressed syllables (Domahs, Wiese, Bornkessel-Schlesewsky, & Schlesewsky, 2008; Knaus, Wiese, & Janben, 2007; Kotz, Schwartze, & Schmidt-Kassow, 2009; Magne, Astesano, Aramaki, Ystad, Kronland-Martinet, & Besson, 2007; Marie, Magne, & Besson, 2011; Quene & Port, 2005; Rothermich, Schmidt-Kassow, & Kotz, 2012) and the syllable rate (Doelling, Arnal, Ghitza, & Poeppel, 2014; Ghitza, 2014; Ghitza & Greenberg, 2009; Shannon, Zeng, Kamath, Wygonski, & Ekelid, 1995) for efficient speech processing. Word stress was also found to facilitate word segmentation at infancy (Dupoux, Pallier, Sebastian, & Mehler, 1997; Höhle, Bijeljac-Babic, Herold, Weissenborn, & Nazzi, 2009; Jusczyk 1997; Shukla, White, & Aslin, 2011) and phrasal level prosody plays an important role in grammar acquisition (Bernard & Gervain, 2012; de Carvalho, Dautriche, Lin, & Christophe, 2017; de Carvalho, Dautriche, & Christophe 2016; Gervain, 2018; Gervain & Werker, 2013). Furthermore, different measures of entrainment to the rhythm of speech was found to be positively associated with speech comprehension and language abilities such as reading skills in adults and children (Abrams, Nicol, Zecker & Kraus, 2009; Ahissar et al., 2001; Giraud & Ramus, 2013).

### 1.2. Rhythm in music and language

Motivated by the highly rhythmical nature of speech, research has explored whether the same mechanisms underlie musical rhythm and speech processing. One of the first theories proposed to account for the shared processes between the domain of music and speech is the Dynamic Attending Theory (e.g., Jones, 2008; Jones & Boltz, 1989; see research on the dynamic nature of attention during the visual processing of stimuli in Busch & VanRullen, 2010 and Dugué, McLelland, Lajous, VanRullen, 2015) emphasizing the role of temporal attention in both domains. The Dynamic Attending Theory states that attention is not equally distributed over time, and for the efficient processing of stimuli with temporal regularities such as music or speech, attention has to be aligned with temporal regularities of the incoming stimuli (e.g., with beats in music or stressed syllables in speech processing). Synchronization of attentional fluctuations with temporal regularities of the auditory input lets the system predict the timing of important events that results in more efficient processing of these stimuli. On the neural level, this was proposed to be achieved by the synchronization or entrainment of neural oscillations with the temporal regularities of incoming stimuli, and several studies support this account both for musical rhythm (e.g., Doelling & Poeppel, 2015; Harding, Sammler, Henry, Large, & Kotz, 2019; Nozaradan, Peretz, Missal, & Mouraux, 2011; Nozaradan, Peretz, & Mouraux, 2012) and speech processing (e.g, Doelling et al., 2014; Ghitza & Greenberg, 2009; Giraud, Kleinschmidt, Poeppel, Lund, Frackowiak, & Laufs, 2007; Harding et al., 2019; Kotz & Schwartze, 2010; Peellee & Davis, 2012; Peelle, Gross, & Davis, 2013).

The role of temporal attention has been primarily emphasized in the processing of surface-level cues in speech (phonemes, word stress) but its role in the processing of hierarchically organized syntactic structures has also been highlighted. The structure of events can be described with hierarchical structures both in rhythm and language (Fitch, 2017; Fitch & Martins, 2014; Lashley, 1951; Lerdahl & Jackendoff, 1983), and initial evidence showing similar electrophysiological responses to violations of rhythmic and linguistic syntax (Sun, Liu, Zhou, & Jiang, 2018) as well as overlapping brain structures involved in hierarchical processing in rhythm and linguistic syntax (Heard & Lee, 2020) indicate shared neural resources underlying syntactic processing in the two domains. Studies showing entrainment of slow oscillations to higher metrical or syntactic boundaries even if these boundaries are not marked by acoustic cues support the role of temporal attention in the processing of syntactic structures in both domains (Nozaradan et al., 2012 for the domain of music and Ding, Melloni, Zhang, Tian, & Poeppel, 2015 for the domain of language).

A number of studies have established associations between various measures of musical rhythmic and phonological (Moritz, Yampolsky, Papadelis, Thomson & Wolf, 2013; Ozernov-Palchik, Wolf, & Patel, 2018; Woodruff Carr et al., 2014) as well as grammatical (Gordon, Jacobs, Schuele, & Mcauley, 2015; Gordon, Shivers, Wieland, Kotz, Yoder, & Mcauley, 2015; Woodruff Carr et al., 2014) skills indicating shared underlying processes for musical rhythm and speech/language processing. In line with these results, populations with language impairments often show weaker rhythmic skills compared to typically developing children. Several studies found an impaired performance on rhythm-related tasks in children with dyslexia (e.g., Muneaux, Ziegler, Truc, Thomson, & Goswami, 2004; Overy, Nicolson, Fawcett, & Clarke, 2003) and with Developmental Language Disorder (Bedoin, Brisseau, Molinier, Roch, & Tillmann, 2016; Corriveau & Goswami, 2009; Corriveau, Pasquini & Goswami, 2007; Cumming, Wilson, Leong, Colling, & Goswami, 2015).

### 1.3. Rhythmic priming

Supporting results for shared neural processes underlying rhythm and language processing motivated an intriguing line of studies that are often referred to as rhythmic priming studies. This work aims to explore whether performance on various linguistic tasks can be improved by boosting the shared network underlying music and language with a presentation of a rhythmic excerpt with a regular beat immediately before the linguistic task. In the majority of the rhythmic priming studies, participants are exposed to an auditory signal (e.g., sequence of percussion beats) with either a regular or irregular beat/non-rhythmic stimuli before completing a linguistic task. Previous research has found an improved performance on a grammaticality judgment task after presentation of a regular rhythmic prime in German-speaking patients with basal ganglia injury (Kotz et al., 2005) and Parkinson disease (Kotz & Gunter, 2015), French-speaking children with dyslexia (Przybylski et al., 2013) and DLD (Bedoin et al., 2016; Przybylski et al., 2013) as well as with typically developing French-speaking (Bedoin et al., 2016; Fiveash, Bedoin, Lalitte, & Tillmann, accepted; Przybylski et al., 2013) and English-speaking (Chern et al., 2018) children. The effect was also found in young adults with and without dyslexia who showed a larger ERP response to grammatical violations following regular vs. the irregular primes (Canette et al., 2020). While most of the studies tested the effect of rhythmic priming on grammaticality judgment, a few works tested its effect on phonological perception in nonexistent words (Cason & Schön, 2012) and sentences (Cason, Astésano, & Schön, 2015). These studies used a slightly different priming paradigm from the grammaticality judgment studies as the rhythm of the primes either matched or did not match the prosodic structure of the speech stimuli. Matching primes facilitated phonological processing compared to not matching primes extending the effect of rhythmic priming to phonemic perception.

The Dynamic Attending Theory offers a potential theoretical account for the rhythmic priming phenomenon. When participants are exposed to music with a regular beat, temporal attention is facilitated by the entrainment of internal oscillators to the beat. As the beat rate of the presented music in the above experiments matches the average rate of stressed syllables in speech (2 Hz), after listening to the musical stimuli the system gets into an ideal state to process speech stimuli. As discussed above, speech also has temporal regularities but in a smaller degree than music, thus without musical stimuli it can take more time for oscillators to synchronize with the regularities of speech stimuli than when musical stimuli are presented before the speech stimuli (see a similar account in Bedoin et al., 2016; Canette et al., 2020; Chern, Tillmann, Vaughan, & Gordon, 2018; Kotz, Gunter, & Wonneberger, 2005; Kotz, Schwartze, & Schmidt-Kassow, 2015; Przybylski et al., 2013). The limited research targeting the neural background of the rhythmic priming effect is in line with the above explanation (Falk, Lanzilotti, & Schön, 2017).

### 1.4. Developmental Language Disorder

In the current paper we investigate the effects of rhythmic priming on language processing and non-linguistic cognitive tasks in typically developing children and children with Developmental Language Disorder (DLD), a developmental disorder formerly known as Specific Language Impairment (SLI) that primarily affects language (with less prominent problems in other domains). Language problems in children with DLD cannot be attributed to obvious impairments in other cognitive domains or perceptual deficits, neurological disorders, emotional or social problems, environmental deprivation or intellectual disability, but the underlying causes of DLD are not well understood yet (Bishop et al., 2017; Leonard, 1998). Traditional accounts suggest a grammar-specific deficit (Clahsen, 1999; Gopnik & Crago, 1991; van der Lely & Stollwerck, 1997; Rice, Wexler & Cleave, 1995), while more recent theories also emphasize the role of domain-general problems and their potential contribution to the linguistic problems (Gathercole & Baddeley, 1990; Hsu & Bishop, 2011; Leonard, 1998; Tallal & Piercy, 1973; Ullman & Pierpont, 2005).

Importantly for our study, several recent studies suggest that rhythm processing is also impaired in DLD (Alcock, Passingham, Watkins & Vargha-Khadem, 2000; Corriveau & Goswami, 2009; Corriveau et al., 2007, Cumming et al., 2015), and this might contribute to the linguistic problems due to the shared underlying mechanisms between rhythm and speech processing discussed above. Weaker rhythm processing in children with DLD is also in line with the Atypical Rhythm Risk Hypothesis that posits that individuals with atypical rhythm are at higher risk for speech/language disorders (Ladányi, Persici, Fiveash, Tillmann & Gordon, 2020).

Goswami (2011) proposed the temporal sampling (oscillatory) model, a theory motivated by the Dynamic Attending Theory (Large & Jones, 1999), to account for language problems primarily in children with dyslexia and, by extension, also for DLD. The framework suggests that inefficient synchronization between brain oscillations and the incoming input contributes to linguistic (and non-linguistic) problems in these populations. Goswami (2011) emphasizes the impairment of processing in the delta band (0.5-3Hz) based on results showing difficulties in tapping to the rhythm at 2 Hz in participants with dyslexia. As the author argues, this is exactly the frequency of occurrence of stressed syllables in English, which explains empirical findings of problems with perceiving syllable stress in dyslexia. Stress is important in language acquisition (Mehler et al., 1988), as well as speech processing, and its impairment can affect all levels of language processing.

### 1.5. The main characteristics of Hungarian

The rhythmic priming effect has so far only been investigated in a few languages (French, English, German), all of which belong to the Indo-European language family and, albeit rhythmically different, have comparable morphosyntactic structure. To test whether rhythmic priming is a language-general phenomenon, it is crucial to test it in languages with various features. In addition, due to its highly complex morphosyntax, Hungarian offers a perfect testing ground to explore whether rhythmic priming has a positive effect when the gap between typical and atypical language development is considerable due to the complex grammar.

Hungarian is an agglutinative language belonging to the Uralic/Finno-Ugric language family. Word order is relatively free, and grammatical relations are mainly expressed by suffixes. Both verbal and nominal morphology is complex. For instance, verbs agree with the number and person of their subject as well as with the definiteness of the object. Tense and mood are also marked. This results in a large number of possible forms. For instance, *olvashatnád*(‘you may/could read it’) consists of the word stem *olvas* (‘read’), the modal suffix*-hat* and the 2^nd^ person singular definite verb suffix in present tense and conditional mood (-*nád*). In addition, both stems and suffixes show strong allophonic variation, e.g., as a result of vowel harmony. Lexical stress in Hungarian is also different from lexical stress in the languages in which rhythmic priming was studied before. Hungarian has fixed stress on the first syllable of words (Siptár & Törkenczy, 2007), which thus provides a cue to word boundaries. The speech rhythm of Hungarian is syllable-timed, similarly to French, but unlike that of English and German. Given these linguistic differences, Hungarian provides a good testing ground to assess the generality of rhythmic priming across languages.

One domain in which Hungarian children with DLD show difficulties is morphology. They make mistakes with case markers (Lukács, Leonard, Kas & Pléh, 2009) and other aspects of verb morphology were also found to be impaired (Leonard, Lukács, & Kas, 2012), as were sentence comprehension and production (Kas & Lukács, 2008). Impairments were found in the lexical domain as well: vocabulary size was shown to be lower (Vinkler & Pléh, 1995) and word retrieval was less efficient (Ladányi & Lukács, 2016) in children with DLD than in their typically developing peers.

### 1.6. The current study

In the current study, we aim to explore three questions: 1) Does the presentation of a regular rhythm improve grammatical processing in Hungarian preschool-aged children with and without Developmental Language Disorder? 2) Is the rhythmic priming effect specific to grammar, general across language domains or general across cognitive domains? 3) Does the rhythmic effect result from the facilitative effect of regular rhythm, the negative effect of the irregular rhythm or both?

Rhythmic priming was shown to be present in a few languages (French, English, German) both in children with and without language disorders. Similarly to the current study, these previous studies also tested the sensitivity to errors in suffixes but those languages have a simpler morphological system than Hungarian. It is possible that exposure to a regular rhythm is sufficient to boost morphological processing in languages with a simple morphology but does not have an effect in morphologically complex languages in which a greater variety of suffixes need to be processed, with sometimes multiple suffixes attached to a stem. For example, in the Chern et al. (2018) study children always had to detect a missing third person singular -*s* as a number agreement error and a missing past tense -*ed* as a tense agreement error. The realization of these error types could not be made more diverse because in English, in the present tense verb conjugation paradigm, all forms are the same (*I/you/we/you/they paint*) except the 3rd person singular ((*s*)*he paints*) and in the past tense paradigm, all forms are exactly the same (*I/you/*(*s*)*he/we/you/they painted*). In contrast, in Hungarian, all six forms are different both for the present and the past tense conjugation paradigm. In addition, the realization of suffixes differs depending on the phonological features of the word stem. To test how the complexity of the Hungarian verb inflection system interacts with rhythmic priming, we included all six forms of the verb conjugation paradigm both for present and past tense suffixes as well as verbs with different phonological features leading to a high diversity in the phonetic realization of the grammatical errors to be detected.

Another feature of Hungarian that distinguishes it from the previously tested languages and which may interact with rhythmic priming is its fixed lexical stress. Hungarian content words are all invariably stressed on their first syllable. This creates a strong rhythmic cue for word segmentation and lexical retrieval. It has never been tested how such a prosodic cue inherent in a language may interact with rhythmic priming.

As discussed above, children with DLD show problems in rhythm processing beyond impairments in language and some other non-linguistic domains. According to the Temporal Sampling Framework (Goswami, 2011), both language and rhythm problems could be partly accounted for by impaired temporal attention due to inefficient synchronization of brain oscillations with the auditory input. It is possible that children with DLD will not show a rhythmic priming effect due to their rhythm processing impairment. Alternatively, the salient and simple beat structure of the rhythmic prime could boost temporal attention, and improve language processing in children with DLD as well.

Previous work in French-speaking children has shown that despite their weaker rhythm skills, children with DLD can also benefit from rhythmic priming. It is, however, unknown whether the effect is also present in a morphologically complex language. It is possible that although children with DLD show a rhythmic priming effect in morphologically simpler languages, but do not do so in morphologically complex languages.

Our first aim is thus to test the generality of rhythmic priming effect in Hungarian-speaking children with and without DLD. Since the relationship between speech and rhythm processing is proposed to be universal as a result of the shared role of temporal attention in rhythm and speech processing across languages, we hypothesize a rhythmic priming effect in Hungarian-speaking children. We expect that rhythmic priming will facilitate grammar processing not only in TD children but also in children with DLD, because processing the rhythm of a regular percussion beat is of greater salience (Ding, Patel, Chen, Butler, Luo, & Poeppel, 2017), lower complexity and higher regularity than processing rhythm in language (Cummins, 2012; Kotz, Ravignani, & Fitch, 2018; Patel, 2008; see also Bedoin et al., 2016; Przybylski et al., 2013). Presentation of regular rhythmic stimuli, therefore, may boost the shared underlying network, presumably through the entrainment of temporal attention to the rhythm of the auditory input, which will then more successfully entrain to speech, leading to more efficient grammatical processing.

Our second aim is to explore whether rhythmic priming is domain-general, specific to language or specific to the processing of spoken language. According to the Dynamic Attending Theory, rhythmic priming appears because internal oscillators entrain to the beat of the regular prime that results in more efficient entrainment with the spoken linguistic stimuli as the beat rate follows the modulation rate of spoken languages. If this mechanism underlies rhythmic priming, it will only work for tasks with stimuli showing such regularity, like speech or rhythm. It is possible, however that rhythmic priming originates from a general arousal in response to the regular prime or from another domain-general mechanism. To test the specificity of rhythmic priming to language processing, we included a non-linguistic task (non-verbal Stroop task) to our design complementing the grammaticality judgment task. Although one previous study (Chern et al., 2018) suggested a language-specific effect, the non-linguistic tasks used in that study differed considerably from the grammaticality judgment task in several respects such as the number of blocks, number of trials within the blocks or the way how trials were presented. In the current study we aimed to match the procedures of our non-grammatical tasks in every possible respect to the grammaticality judgment task. Furthermore, similarly to Chern et al. (2018)’s study, we chose a task that requires attentional processes but stimuli do not involve regularities as speech or rhythm does.

In addition to the non-linguistic task, we also included a linguistic but non-grammatical task to our design to examine the possibility that rhythmic priming is not domain-general but is also present in language production beyond language processing. Underlying brain networks for temporal processing in speech perception and production has been proposed to highly overlap (Kotz & Schwartze, 2010) and this network was proposed to underlie temporal processing of musical rhythm (Kotz et al., 2018). If rhythmic priming boosts overlapping neural functions underlying rhythm perception and speech production, then rhythmic priming could also appear in a picture naming task. If the rhythmic priming effect appears a result of facilitation of temporal attention, it will not be present in the picture naming task. In line with the Dynamic Attending Theory, we expect to find it only in the grammaticality judgment task and not in the picture-naming task or the non-verbal Stroop task.

Our third aim is to better understand the origin of the rhythmic priming effect by comparing performance in three within-subject conditions: one in which a regular rhythm is used as a prime, one with an irregular rhythm and a baseline condition in which children were not presented with any auditory stimuli. Previous studies compared performance only between two conditions (regular vs. irregular: Chern et al., 2018; Przybylski et al., 2013; regular vs. nonrhythmic baseline condition: Bedoin et al., 2016), which did not allow for a direct comparison of the irregular prime to a non-rhythmic baseline condition. A better performance following a regular prime compared to an irregular prime can appear 1) due to the synchronization of oscillations with the regular rhythm that would facilitate subsequent spoken language processing with a similar modulation rate to the rhythm, 2) due to the alignment of oscillations to the irregular rhythm that would hinder spoken language processing or 3) both of these mechanisms. The result on better grammaticality judgment performance after a regular rhythm compared to a non-rhythmic environmental noise (Bedoin et al., 2016) excludes the possibility that rhythmic priming results only from the negative effect of the irregular prime (Explanation 2). It is still not known, however, whether rhythmic can be accounted for by only the positive effect of the regular prime (Explanation 1) or the combination of the positive effect of the regular prime and the negative effect of the irregular prime (Explanation 3). By including three conditions, we will be able to decide between these two options, and better understand mechanisms underlying rhythmic priming.

We expect to find a better performance after the regular prime than after the irregular prime and the baseline condition as a result of the facilitating effect of regular primes on temporal attention and the lack of such effect in the case of the irregular prime and the baseline condition. As Bedoin et al. (2016) discusses, the effect size of the difference between the regular and the irregular condition was larger in Przybylski et al.’s (2013) study than between the regular and the non-rhythmic baseline conditions in their study, suggesting that the irregular prime may have a negative effect on grammatical processing. Based on this observation, we expect to find weaker performance in the irregular condition than in the baseline condition.

## 2. Methods

### 2.1. Participants

Seventeen 6-year-old children with DLD and 17 age- and IQ-matched typically developing children participated in the study. Demographic and screening data for the two groups are shown in Table 1. Children were recruited from six different pre-schools in Budapest. As a first step of recruitment, a larger group of children showing language problems was selected by the speech-language therapists of the pre-schools. Children whose parents agreed to participate in the study (n = 50) were screened for DLD by a speech-language therapist using four language tasks (the Hungarian version of the Peabody Picture Vocabulary Test (Csányi, 1974), the Hungarian adaptation of the Test for Reception of Grammar (Bishop, 1983, 2012; Lukács, Győri, & Rózsa, 2012), the Hungarian Sentence Repetition Test (Magyar Mondatutánmondási Teszt; Kas & Lukács, in preparation), and a nonword repetition test (Racsmány, Lukács, Németh, & Pléh, 2005) as well as a non-verbal intelligence task (Raven Colored Progressive Matrices, Raven, Court, & Raven, 1987)). Children who scored at least 1.5 SDs below age norms on at least two out of the four language tasks and had an IQ above 85 were selected for the DLD group (n = 17). After selecting the DLD group, typically developing children matching in age and gender to children in the DLD group were selected by the teachers in the same pre-schools. Children whose parents agreed to participate in the study completed the non-verbal intelligence task. One TD child matching in IQ (< 5 points of difference in IQ), age (< 3 months difference) and sex was selected for each DLD child. We made every effort to match the groups in parent education level, but this aim could only partially be met, and the educational level of the DLD group is slightly lower than that of the TD group (see Table 1). TD children also completed the screening tasks to make sure that they don’t have language impairment. None of the TD children performed below average on the screening tasks. All children were tested with the informed consent of their parents, in accordance with the principles set out in the Declaration of Helsinki and the stipulations of the local Institutional Review Board.

**Table 1.** Demographic data and results on the screening tests in typically developing children and children with DLD (PPVT – Peabody Picture Vocabulary Test, TROG – Test for Reception of Grammar)

|  | **TD mean (SD)** | **DLD mean (SD)** | **difference** |
| --- | --- | --- | --- |
| **age** | 74 mo (6 mo) | 74 mo (6 mo) | $t(16) = 1.201, n.s.$ |
| **IQ (Raven)** | 104 (10) | 102 y (9) | $t(16) = .890, n.s.$ |
| **Parental education** | 14 y (1.8 y) | 12 y (1.6 y) | $t(16) = 3.267 p = .005$ |
| **PPVT** | 82 (17) | 63 (8) | $t(16) = 4.258 p = .001$ |
| **TROG** | 65 (4) | 50 (11) | $t(16) = 6.520 p < .001$ |
| **sentence repetition** | 30 (9) | 20 (9) | $t(16) = 3.154 p = .006$ |
| **non-word repetition** | 4.7 (0.9) | 2.8 (1.2) | $t(16) = 5.343 p < .001$ |

### 2.2. Tasks

#### 2.2.1. Grammaticality judgment task

##### Stimuli

We created 60 grammatically correct Hungarian sentences. Then we modified these sentences by violating either the number agreement between the verb and the subject or the tense of the verb with respect to the adverb (Table 2). All 6 forms (singular and plural forms, in 1^st^, 2^nd^, and 3^rd^ person) of the present and past verb conjugation paradigm were equally represented both in grammatical and ungrammatical sentences. Two lists of sentences were created by combining 30 grammatically correct and 30 grammatically incorrect sentences. We included correct and incorrect versions of the same sentence in different lists, therefore one child couldn’t hear both (see examples in Table 2), and lists were randomly assigned to children. We also made sure that the two lists were matched in their sentence types and vocabulary. Sentences were recorded by an adult female speaker in a mild child-directed manner.

**Table 2.** Examples of grammatically correct and incorrect sentences in the rhythmic priming grammaticality judgment task

|  | Correct version | Violation |
| --- | --- | --- |
| Number agreement | *A kisbaba alszik az ágyban.*<br>'The baby is sleeping in the bed.' | *\*A kisbaba alszanak az ágyban.*<br>'The baby are sleeping in the bed.' |
| Tense | *Tegnap este sok fagyit kaptunk.*<br>'We got a lot of ice-cream last evening' | *\*Tegnap este sok fagyit kapunk.*<br>'We get a lot of ice-cream last evening' |

As regular and irregular primes, we used the same stimuli as several previous studies (Bedoin et al., 2016; Chern et al., 2018; Przybylski et al., 2013). Both sequences were played on a tam-tam and maracas and contained the same number of tones. The regular prime had a rhythmic structure from which it was relatively easy to extract the meter, while it was more difficult to extract the rhythmic structure from the irregular prime (see a more detailed description of the primes in Przybylski et al., 2013).

##### Design and Procedure

During the grammaticality judgment task, children were presented with six blocks of ten sentences on which they were asked to make grammaticality judgments (see a summary of the task procedures in Figure 1). Each block was preceded by a 32-second-long prime. Two primes belonged to the regular condition, two belonged to the irregular condition and two primes belonged to the baseline condition, which did not contain any auditory stimuli. Prime conditions followed each other in a pseudo-randomized order with the restriction that two primes from the same condition could not follow each other. A picture of a cat relaxing and listening to music was visually presented on the screen while the prime stimuli were played. Children were instructed to relax and take some rest during the primes.

**Figure 1.**
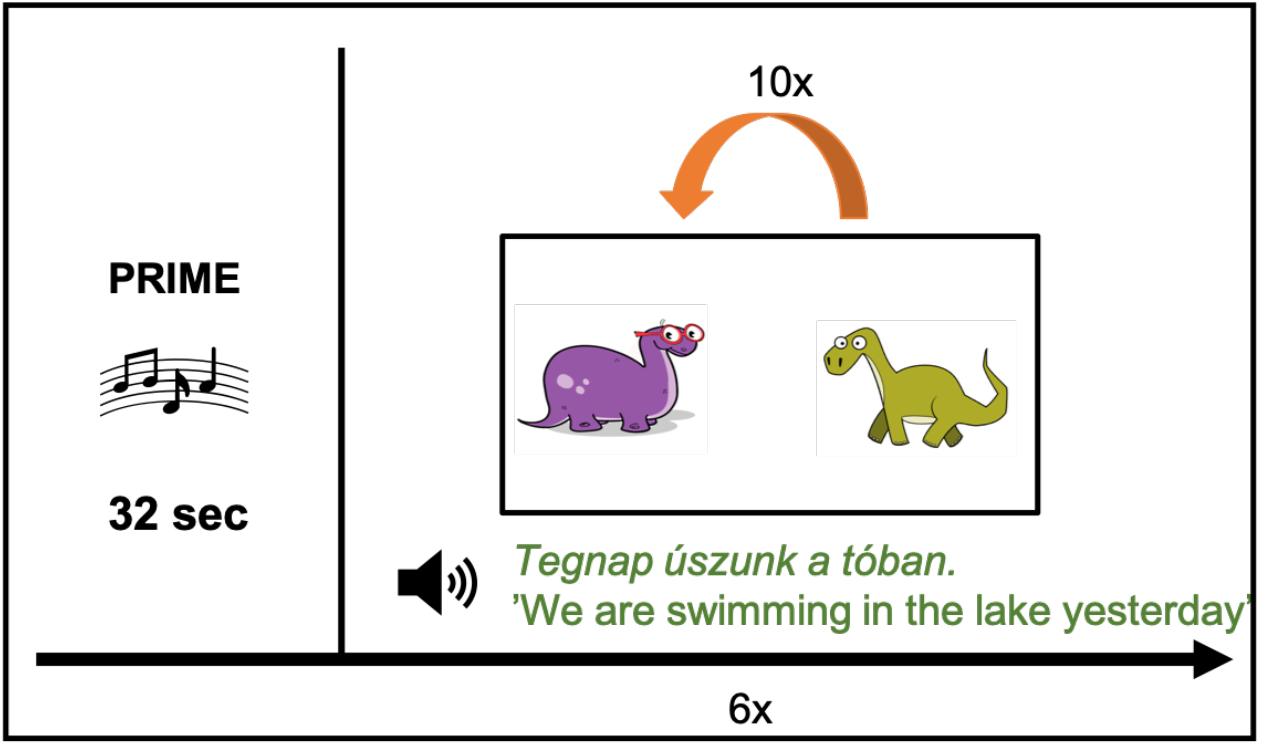
Procedures of the grammaticality judgment rhythmic priming task. The grammaticality judgment task consisted of 6 blocks. In each block, first, children were presented with a prime, and then with 10 sentences half of which was grammatical and the other half was ungrammatical.

Each block contained five grammatically correct and five grammatically incorrect sentences, which were presented to the child in a randomized order. The type of violation was counterbalanced across blocks.

We created a background story similar to the one used in previous studies (Bedoin et al., 2016; Chern et al., 2018; Przybylski et al., 2012) to make our task more comprehensible and interesting for young children. Before the task, we introduced them a purple and a green dinosaur. They were informed that the purple dinosaur – which looked very confident and attentive – was very attentive, so it said everything correctly, while the green dinosaur – which looked somewhat confused – was less attentive, and always made mistakes. We presented them with examples of the sentences the dinosaurs produced. As the first part of the practice phase, children heard the same sentence, once in a correct form, once in an incorrect form, and they had to decide which dinosaur produced the sentence. A second practice phase followed, in which children were presented with different correct and incorrect sentences and they had to decide which dinosaur produced the sentence. Children got feedback for each response during the practice phase. After we made sure that children understood the task, we moved on to the test.

We used d’ scores to measure grammaticality judgment performance, similarly to previous studies (Bedoin et al., 2016; Chern et al., 2018; Przybylski et al., 2013). D’ is a measure used by signal detection theory that describes the participant’s ability to discriminate between two types of stimuli (in the current study between grammatical and ungrammatical stimuli) and was calculated as the difference between the z-transformed hit rate and false alarm rate (d’ = z(hit rate)-z(false alarm rate).

We presented stimuli and collected data with OpenSesame (Mathôt, Schreij, & Theeuwes, 2012) for all the three rhythmic priming tasks.

#### 2.2.3. Picture naming task

##### Stimuli

Sixty pictures of simple everyday objects that the children likely knew were selected from the picture set used by the norming study of Bates and colleagues (2003). Pictures were grouped into six blocks. The same auditory stimuli were used as primes as in the grammaticality judgment task.

##### Design and procedure

To make sure that children were familiar with the names of the pictures, they were asked to name all the pictures in a randomized order at a previous testing session.

The structure of the picture naming task was the same as the structure of the grammaticality judgment task. Children were presented with a prime and then named 10 pictures. Children were asked to name pictures as quickly and as accurately as possible. Pictures were presented in a randomized order within a block and blocks of pictures were randomly assigned to a prime condition. Children’s responses were audio-recorded. Naming errors and response times of the correct responses were coded based on the recording. We decided to code naming times off-line for a more accurate measure. Based on our experience with naming studies with young children, automatic measurements detecting the onset of the name are problematic, since children often make false starts, emitting sounds other than the name before actually naming the picture, which activates the voice key. Children also often produce or start to produce names too softly, which fails to activate the voice key. Both of these lead to inaccurate measures (see also Cummings, Seddoh, & Jallo, 2016). We excluded incorrect answers and answers with naming times above 5 seconds, since they indicate that the child was inattentive at that trial, and calculated mean naming times for the three conditions. Accuracy was also coded and calculated, but since performance was at ceiling, we did not analyze accuracy data statistically.

#### 2.2.4. Non-verbal Stroop task

##### Stimuli

For the non-verbal Stroop task, we created arrows appearing on a blank screen. Arrows pointed to the left, to the right, up or down and appeared either in the middle of the screen (baseline condition) or in the left, right, top or bottom of the screen either in a congruent position with the direction of the arrow (e.g., an arrow pointing up appears at the top of the screen) or in an incongruent position with the direction of the arrow (e.g., an arrow pointing up appears at the bottom of the screen). Six blocks of 20 trials were created. The number of arrows with a congruent, incongruent and middle position were counterbalanced across blocks as well as the number of arrows pointing to right, left, up and down.

As primes, we used the same auditory stimuli as in the grammaticality judgment task and the picture naming task.

##### Design and procedure

The structure of the task was the same as in the grammaticality judgment task and the picture naming task. Children were presented with a prime and then with a block of 20 arrows. Children were asked to press a button based on the direction of the arrow. Arrows followed each other in a randomized order in each block and associations between arrow blocks and prime conditions were also randomized. The task was preceded by a practice session. We calculated two different measures for the non-verbal Stroop task. First, for testing the effect of rhythmic priming on executive functions, we calculated mean Stroop effects measured by accuracy and reaction times for the three priming conditions. Stroop effects were calculated as the difference between mean reaction times for correct responses/accuracy for the incongruent and baseline conditions. Second, for testing the effect of rhythmic priming on the general performance on the task, accuracy and mean reaction times were calculated with collapsing across Stroop conditions for the three priming conditions. Trials with reaction times under 300 ms-s and above 5000 ms-s were excluded from the analysis as these data resulted from technical errors or inattentiveness of the child, and are not meaningful.

## 3. Results

### 3.1. Grammaticality judgment task

Performance on the grammaticality judgment task is shown in Table 3 and Figure 2. We conducted a 2 x 3 ANOVA with Group as a between-subject factor (DLD vs. TD) and Rhythm (Regular vs. Irregular vs. Baseline) as a within-subject factor.

**Figure 2.**
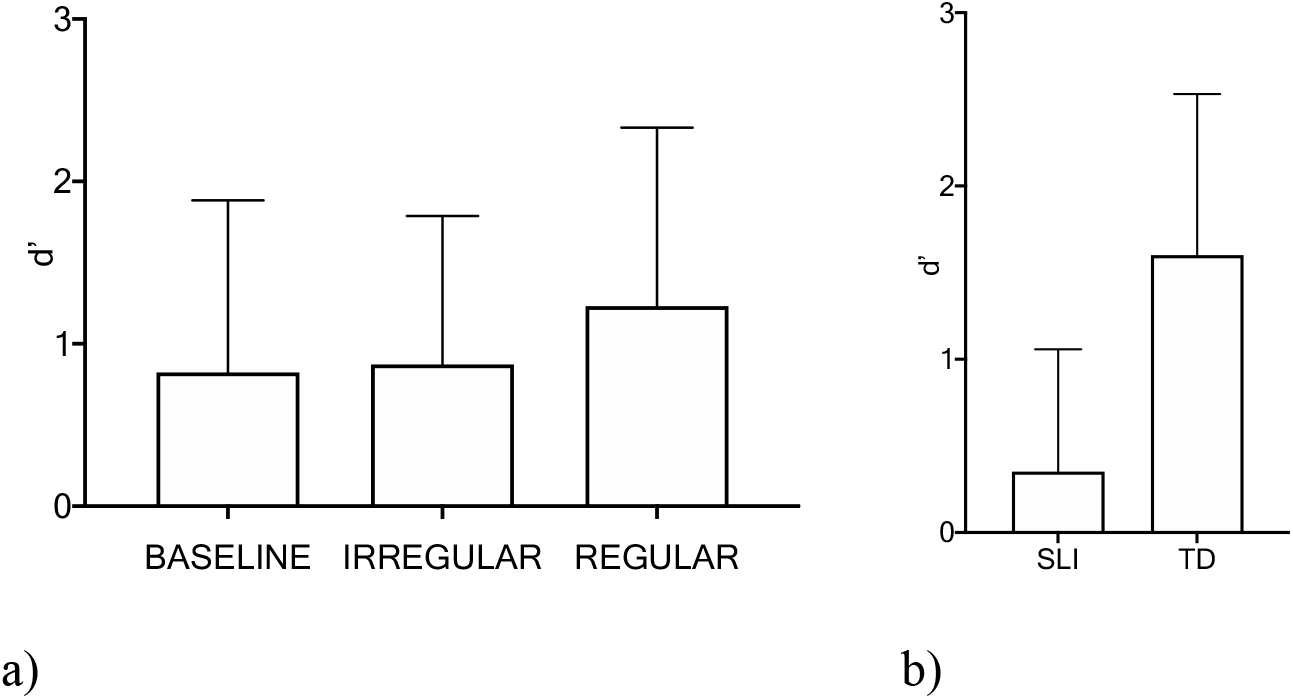
a) Performance on the grammaticality judgment task in the three rhythm conditions collapsed across TD and DLD groups. b) Group difference in the grammaticality judgment task. Error bars represent standard deviations.

**Table 3a-b.** a) Means and standard deviations of hit rates, false alarm rates in the DLD and TD groups in the three rhythm conditions. b) Means and standard deviations of z-scored hit rates, false alarm rates and d’ scores in the DLD and TD groups in the three rhythm conditions.

|  | Hit rate |  |  | False alarm rate |  |  |
| --- | --- | --- | --- | --- | --- | --- |
|  | Baseline | Irregular | Regular | Baseline | Irregular | Regular |
| **DLD** | 0.61<br>(0.24) | 0.63<br>(0.23) | 0.68<br>(0.22) | 0.57<br>(0.20) | 0.52<br>(0.27) | 0.52<br>(0.27) |
| **TD** | 0.81<br>(0.14) | 0.77<br>(0.16) | 0.77<br>(0.17) | 0.32<br>(0.20) | 0.30<br>(0.14) | 0.25<br>(0.21) |

| | Hit (z-score) | | | False alarm (z-score) | | | $d'$ | | |
| --- | --- | --- | --- | --- | --- | --- | --- | --- | --- |
|  | Baseline | Irregular | Regular | Irregular | Regular | Regular | Baseline | Irregular | Regular |
| **DLD** | 0.36<br>(0.83) | 0.42<br>(0.80) | 0.63<br>(0.85) | 0.25<br>(0.65) | 0.10<br>(0.80) | 0.01<br>(0.83) | 0.12<br>(0.76) | 0.32<br>(0.73) | 0.62<br>(0.88) |
| **TD** | 0.97<br>(0.56) | 0.86<br>(0.61) | 0.90<br>(0.70) | -0.56<br>(0.70) | -0.57<br>(0.43) | -0.94<br>(0.91) | 1.53<br>(0.90) | 1.43<br>(0.73) | 1.84<br>(1.13) |

The main effect of Group was significant (*F*(1, 32) = 42.41, p < .001) showing a significantly higher performance in children with TD (mean *d’* = 1.60, *SD* = 0.93) than in children with DLD (mean *d’* = 0.35, *SD* = 0.71). The main effect of rhythm was also significant (*F*(1, 32) = 3.15, *p* = .0496). Fisher’s Least Significant Difference (LSD) post-hoc pairwise comparisons showed a significantly better performance in the regular (mean *d’* = 1.23, *SD* = 1.10) than in the baseline (mean *d’* = 0.82, *SD* = 1.06; *t*(33) = 2.117, *p* = .025, *d* = 0.38) and irregular (mean *d’* = 0.873, *SD* = 0.91; *t*(33) = 2.00, *p* = .047, *d* = 0.36) conditions. There was no significant difference between the irregular and baseline conditions (*t*(33) = 0.327, *p* = .779, *d* = .05). The Group X Rhythm interaction was not significant. To confirm the reliability of the significant effect of rhythm on grammaticality judgment, we also ran a Bayesian repeated measures ANOVA. This analysis also showed that the model with the main effect of rhythm and DLD is the best model of the data. The probability of the observed data was 42594 times greater with this model than the null model.

Since the DLD group showed a weak performance on the grammaticality judgment task, we tested whether d-prime scores on different conditions significantly differ from chance level (*d’* = 0) with a one-sample t-test in the TD and DLD groups, separately. We found that while children with TD showed a performance significantly above chance in each condition (baseline: *t*(16) = 7.012, *p* < .001, irregular = *t*(16) = 8.1, p < .001, regular: *t*(16) = .6.745, *p* < .001), children with DLD performed above chance only in the regular rhythm condition (baseline: *t*(16) = .715, *p* = .485, irregular *t*(16) = 1.786, *p* = .093, regular: *t*(16) = 3.897, *p* = .001).

As Bedoin et al. (2016) argue, using silence as the baseline condition runs a risk that the effect of the previous prime persists and affects the performance in the silent baseline condition. To test this possibility, we grouped the responses of the baseline condition into three categories: (i) baseline block preceded by a regular block, (ii) baseline block preceded by an irregular block and (iii) baseline as the first block in the task. We calculated the mean d’ scores for each category. We did not find a significant difference between these cases in a one-way ANOVA (*F*(2, 64) = 1.452, *p* = .242; *M_irregular_* = 0.97 *M_regular_* = 0.78 *M_first_* = 2.06). To investigate the possibility that the prime from the previous block only affected the first items of the block, we ran the same analyses only on the first five items from each block but no significant difference appeared between blocks in this analysis either (*F*(2, 64) = 0.371, *p* = .692; *M_irregular_* = 1.46 *M_regular_* = 0.94 *M_first_* = 1.15).

### 3.2. Picture naming task

Mean naming times in the picture naming task are shown in Table 4. Naming times for the correct answers were analyzed in a 2 x 3 ANOVA with Group (DLD vs. TD) as a between-subject factor and Rhythm (Regular vs. Irregular vs. Baseline) as a within-subject factor. The main effect of Group was marginally significant (*F*(1, 32) = 4.14, *p* = .050) reflecting slower naming times in the DLD (*M* = 1404 ms, *SD* = 422 ms) than in the TD group (*M* = 1177 ms, *SD* = 370 ms). There were no other significant effects or interactions.

**Table 4.** Mean naming times and standard deviations in milliseconds in the picture naming task

|  | **Baseline** | **Irregular** | **Regular** |
| --- | --- | --- | --- |
| **DLD** | 1450 (109) | 1328 (86) | 1435 (114) |
| **TD** | 1183 (101) | 1156 (91) | 1192 (81) |

### 3.3. Non-verbal Stroop task

Differences in Stroop effects depending on the priming condition were analyzed both measured by accuracy and reaction times as the dependent variable with 2 x 3 ANOVA-s with Group (DLD vs. TD) as a between-subject factor and Rhythm (Regular vs. Irregular vs. Baseline) as a within-subject factor (see a summary of Stroop scores in Table 5). The interaction between Group and Rhythm was significant in the case of reaction times (*F*(2, 64) = 4.23, *p* = .019). Fisher’s Least Significant Difference (LSD) post-hoc pairwise comparisons showed a significantly smaller Stroop effect in the case of the irregular than in the baseline (*t*(16) = 2.53, *p* = .010) and in the regular (*t*(16) = 2.53, *p* = .020) condition in the TD group but no significant differences appeared in the DLD group. No other main effects or interactions were significant in the case of Stroop effects measured by reaction times or accuracy rates.

**Table 5.** Mean Stroop effects (difference between mean reaction times for the baseline and incongruent conditions) in the non-verbal Stroop tasks. Reaction time data are reported in milliseconds.

|  | Mean Stroop effect reaction time |  |  | Mean Stroop effect accuracy rate |  |  |
| --- | --- | --- | --- | --- | --- | --- |
|  | Baseline | Irregular | Regular | Baseline | Irregular | Regular |
| **DLD** | 213 (282) | 324 (231) | 288 (345) | 0.04 (0.13) | 0.11 (0.10) | 0.07 (0.08) |
| **TD** | 307 (240) | 91 (277) | 286 (283) | 0.08 (0.14) | 0.08 (0.12) | 0.09 (0.17) |

To explore whether rhythm had an effect on task performance in general, independently of the Stroop effect, accuracy and reaction times of correct responses collapsed across Stroop conditions were analyzed with 2 x 3 ANOVA-s with Group (DLD vs. TD) as a between-subject factor and Rhythm (Regular vs. Irregular vs. Baseline) as a within-subject factor (see a summary of scores in Table 6). No main effects or interactions were found to be significant either for accuracy, or for reaction times.

**Table 6.**
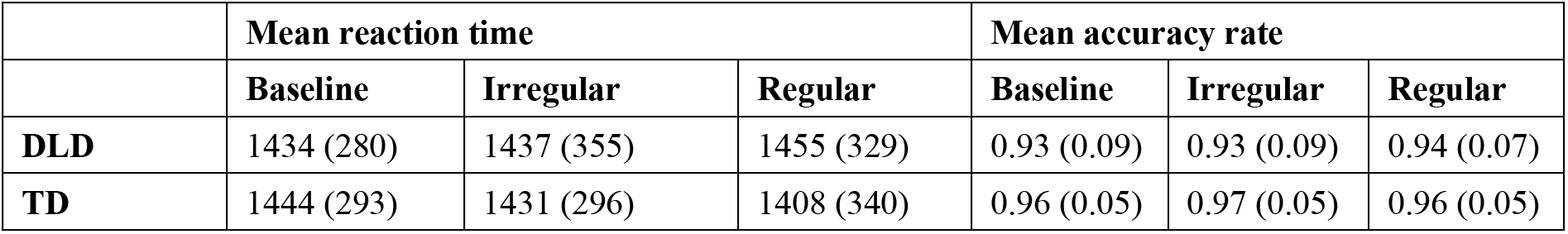
Mean reaction times and standard deviations in the non-verbal Stroop task. Reaction time data are reported in milliseconds.

## 4. Discussion

In the current study, we aimed to investigate the rhythmic priming effect in Hungarian-speaking 5-7-year-old children with and without Developmental Language Disorder. Following exposure to a regular vs. an irregular rhythm or no prime, children showed a rhythmic priming effect, i.e. a higher performance, on a grammaticality judgment task, but did not on a picture naming task and a non-verbal Stroop task, requiring no grammatical processing. The results support the prediction that a rhythm with a regular metrical structure facilitates subsequent grammatical processing in an agglutinating language, a morphosyntactic type never tested before. This is in line with the hypothesis that facilitation of entrainment of internal with regularities of the auditory input underlies rhythmic priming. The presence of the effect did not depend on the presence or absence of DLD. Children with DLD also benefitted from the regular rhythmic prime, suggesting that rhythmic training or the use of rhythmic stimuli during language therapy might facilitate language processing in this atypical group. This finding has important implications for speech-language therapy for DLD.

Our finding about the presence of the rhythmic priming effect in a grammaticality judgment task in children with and without DLD is in line with previous studies finding better performance when children are exposed to a regular rhythm compared to an irregular one (Chern et al., 2018; Przybylski et al., 2013) or no rhythm at all (Bedoin et al., 2016). Our study also extends previous research by showing evidence for a longer effect of priming than previous studies as in the current study, children were presented with 10 sentences after the prime in the grammaticality judgment task that is slightly more than in previous studies in that 6 sentences were presented per block.

Regarding the difference between the baseline and irregular conditions in the grammaticality judgment task, we expected to find a weaker performance in the irregular condition than in the baseline condition based on Bedoin et al.’s (2016) observation about the rhythmic priming effect being stronger when comparing the regular condition to an irregular condition than when comparing the regular condition to a non-rhythmic baseline. In the current study, we were able to directly compare the irregular and baseline conditions and found no significant difference between them. This suggests that irregular rhythm does not have a negative effect on subsequent grammatical processing. This pattern of results indicates that while rhythm with a regular beat can facilitate temporal attention during the processing of subsequently presented spoken linguistic stimuli, rhythm with an irregular beat does not lead to a disruption of temporal attention in subsequent language processing. We note, however, that while Bedoin et al. (2016) used environmental noise as a baseline condition, we used silence; perhaps environmental noise and silence impact performance differently.

The presence of the rhythmic priming effect in the grammaticality judgment task and its absence in the non-verbal Stroop task, a non-linguistic task strongly relying on attentional processes, is in line with previous results (Chern et al., 2018). In the current study, the length of the blocks as well as the procedures of the task were closely matched between the three tasks. These results suggest that the rhythmic priming effect is specific to speech/language processing or to grammar processing. This is in line with the temporal attention account of the rhythmic priming effect as stimuli did not appear in a temporarily regular fashion in the case of the nonverbal Stroop task in contrast to the regularities of speech stimuli in the grammaticality judgment task. Note that in the non-verbal Stroop task, children with TD showed a reduced Stroop effect in the irregular condition compared to the baseline or regular conditions. This result could be explained by the fact that the Stroop effect in the irregular condition was negative in seven out of the seventeen children with TD meaning that it took longer for them to respond to baseline trials than to Stroop trials. Missing Stroop effects in the irregular condition in some children are difficult to interpret but we believe that this result is unrelated to rhythmic priming, therefore we do not discuss it further.

To our knowledge, our study is the first one exploring the effect of rhythmic priming on word retrieval. The lack of rhythmic priming in the picture naming task suggests that the rhythmic priming effect is not only specific to the language domain but seems to be restricted to language perception or grammar. These results are in line with a recent study (Canette, Lalitte, Bedoin, Pineau, Bigand, & Tillmann, 2020) testing the effect of a rhythmical music vs. textural sounds on a semantic evocation task in which children were asked to verbalize concepts that were evoked by rhythmic music or sound texture. The study found that children produced more concepts after the textural sounds suggesting that rhythmic priming did not facilitate semantic evocation. Although picture naming and semantic evocation measure partly different abilities, both involve lexical processes. The lack of rhythmic priming in lexical tasks further confirms that rhythmic priming is specific to grammatical or language processing, and supports the interpretation that facilitation of temporal attention accounts for the rhythmic priming effect. Alternatively, it is possible that rhythmic priming could facilitate speech processing in other tasks focusing on the production of longer units, such as sentences in that the temporal “grid” (see Kotz & Schwartze, 2016) of the sentence is longer than for single words or in patients with impaired temporal processing such as individuals with basal ganglia lesions or Parkinson disease.

The design of the current study did not enable us to decide which aspect of language processing – lower-level auditory processing of phonemes and syllables, the processing of hierarchical linguistic structures or both – is facilitated by the regular prime, neither to explore the underlying mechanisms. The result that rhythmic priming facilitates phonological processing (Cason, Astésano, & Schön, 2015; Cason & Schön, 2012) supports the role of auditory processing, but future studies need to find out whether improved grammatical processing is accounted for by better auditory processing in itself or rhythmic priming facilitates both auditory processing and grammatical processing.

The mechanisms underlying rhythmic priming are also still to be discovered. The Dynamic Attending Theory offers an explanation that could account for the facilitative effect of rhythm both on the processing of surface-level auditory cues and grammatical processing. Oscillators synchronized with surface-level cues would facilitate auditory processing, while slower oscillators synchronized with syntactic units would facilitate syntactic processing. According to another explanation, facilitation of temporal attention affects sequencing in speech perception, and that would lead to the more efficient detection of errors (see Bedoin et al., 2016; Kotz & Schwartze, 2009; Przybylski et al., 2013).

It is also important to emphasize that a grammaticality judgment task requires strong metalinguistic skills that makes it a less direct measure of grammatical processing. While good grammatical skills are obviously necessary to detect grammatical errors, they are not sufficient. As language processes are highly automatic, it is possible that one has good grammatical skills and is able to process sentences but is less successful at detecting errors that requires metalinguistic skills and attention to violations. In this case, their performance will show a high hit rate with a relatively high false alarm. High false alarms played a stronger role in low d’-s than low hit rates, especially in the DLD group, meaning that judging ungrammatical sentences “correct” accounted for more errors than judging grammatical sentences “incorrect”. This could mean that metalinguistic skills, attentional problems or the usual “yes”-bias commonly found in children also played a role in the performance on the grammaticality judgment task. Future studies will need to investigate the effect of rhythmic priming on grammatical processing using behavioral tasks or EEG measuring on-line grammatical processing. High false alarms played a stronger role in low *d’*-s than low hit rates, especially in the DLD group, meaning that judging ungrammatical sentences “correct” accounted for more errors than judging grammatical sentences “incorrect”. This could mean that metalinguistic skills or attentional problems also played a role in the performance on the grammaticality judgment task, and potentially were affected by rhythmic priming. Future studies will need to investigate the effect of rhythmic priming on grammatical processing using behavioral tasks or EEG measuring on-line grammatical processing.

Finding the mechanisms underlying rhythmic priming could also help design ways to use rhythm in general (see also Schön & Tillman, 2015), or rhythmic priming in particular (Bedoin et al., 2017) efficiently in speech-language therapy. Further rhythmic studies with manipulating the rhythm and the target task and measuring brain activity during both the rhythmic priming stimuli and the presentation of the linguistic stimuli will help find the sources of the effect. If future research will support the efficiency of the use of rhythmic stimuli during speech-language therapy, then rhythmic training or presentation of rhythmic stimuli during tasks targeting speech-language skills should be included in therapy.

## Acknowledgements

This study was conducted with the support of the European Union’s Horizon 2020 research and innovation program under the Marie Sklodowska-Curie grant agreement No. 641858 (PredictAble project) to J.G and E.L, the ERC Consolidator Grant (773202 ERC-2017-COG ‘BabyRhythm’; https://erc.europa.eu/funding/consolidator-grants) to J.G. Á.L. was supported by the Hungarian Academy of Sciences Lendület Program (96233 MTA-BME Lendület Language Acquisition Research Group). E.L. was also supported by funding from the National Institutes of Health: the NIH Common Fund through the Office of NIH Director, and the National Institute on Deafness and Other Communication Disorders, under Award Numbers DP2HD098859 and R01DC016977. The content is solely the responsibility of the authors and does not necessarily represent the official views of the NIH. The authors declare no conflicts of interest with regard to the funding source for this study.

Special thanks go to the children, their teachers and parents for their kindness and cooperation. We thank Ágnes Kovács for her inevitable help in screening our participants and Lilla Zakariás for her help in recording the stimuli and helpful advices about task procedures. A huge thank goes to Dr. Reyna Gordon and the Music Cognition Lab of Vanderbilt University Medical Center, Nashville for the fruitful discussions and motivating environment.

## Data Availability Statement

The data that support the findings of this study are available from the corresponding author upon reasonable request from Eniko Ladanyi at the following email address.

